# Phenological displacement is uncommon among sympatric angiosperms

**DOI:** 10.1101/2020.08.04.236935

**Authors:** Daniel S. Park, Ian K. Breckheimer, Aaron M. Ellison, Goia M. Lyra, Charles C. Davis

## Abstract

Interactions between species can influence access to resources and successful reproduction. One possible outcome of such interactions is reproductive character displacement. Here, the similarity of reproductive traits – such as flowering time – among close relatives growing in sympatry differ more so than when growing apart. However, evidence for the overall prevalence and direction of this phenomenon, or the stability of such differences under environmental change, remains untested across large taxonomic and spatial scales. We apply data from tens of thousands of herbarium specimens to examine character displacement in flowering time across 110 animal-pollinated angiosperm species in the eastern USA. We demonstrate that the degree and direction of phenological displacement among co-occurring closely related species pairs varies tremendously. Overall, flowering time displacement in sympatry is not common. However, displacement is generally greater among species pairs that flower close in time, regardless of direction. We additionally identify that future climate change may alter the nature of phenological displacement among many of these species pairs. On average, flowering times of closely related species were predicted to shift further apart by the mid-21st century, which may have significant future consequences for species interactions and gene flow.

## Introduction

Interactions between species can affect access to resources and successful reproduction. The outcome of such interactions may result in character displacement, in which the phenotypic similarity of species differs depending on whether they are co-occurring (sympatry) or not (W. L. Brown & Wilson, 1956; Connell, 1980; Grant, 1972). Numerous instances of character displacement have been identified across the tree of life (Dayan & Simberloff, 2005; Pfennig & Pfennig, 2009). However, evidence for the overall prevalence and direction of this phenomenon or the stability of such differences under future environmental change is lacking (Hopkins, 2013; Levin, 2006).

Reproductive character displacement – the modification of reproductive traits in sympatric populations of related species – is widely considered to be a key mechanism facilitating co-occurrence, reproductive isolation, and ecological and evolutionary divergence (J A Coyne & Orr, 2004; Jerry A Coyne, 1992; Grant & Grant, 2011; Mayr, 1947). This is especially true for the timing (phenology) of flowering, which is strongly linked to fitness and often highly variable even among closely related taxa (Briscoe Runquist, Chu, Iverson, Kopp, & Moeller, 2014; Brody, 1997; Domínguez & Dirzo, 1995; Galloway, 2002; Kelly & Levin, 2000; Lacey, Roach, Herr, Kincaid, & Perrott, 2003; Lowry, Rockwood, & Willis, 2008; Park et al., 2018; B Rathcke & Lacey, 1985; Sletvold, Moritz, & Ågren, 2015; Spriggs et al., 2019; Stinson, 2004). Plants often flower and share pollinators with other species across their range, and this community context has been demonstrated to greatly influence reproductive phenology (B. J. Brown, Mitchell, & Graham, 2002; Moeller, 2004; Stiles, 1975, 1977).

Flowering phenology is a heritable trait on which selection can act rapidly (Allard & Hansche, 1964; Bergh, 1976; Izawa, 2007). Despite its relevance, empirical evidence for phenological character displacement in plants remains limited to a small number of case studies (e.g., (Briscoe Runquist et al., 2014; Lowry et al., 2008; Spriggs et al., 2019). This greatly limits our ability to understand the general relevance of phenological displacement governing plant interactions and distributions. Moreover, flowering phenology is highly responsive to climate (Davis, Willis, Connolly, Kelly, & Ellison, 2015; Franks, Sim, & Weis, 2007; Sherry et al., 2007), and it remains an open question whether current phenological similarities or differences among co-occurring species are likely to remain constant in the face of future climate change.

Phenological character displacement is commonly inferred to imply phenological divergence in sympatry, but it can also manifest as phenological convergence; the nature of interspecific interactions will determine which applies (Grant, 1972). For example, two species may *diverge* in flowering time when they co-occur (Fig. 1a), thus reducing competition (Campbell, 1985; Elzinga et al., 2007; Stone, Willmer, & Rowe, 1998). Such asynchronous flowering also can reproductively isolate species and reduce the costs of heterospecific pollen transfer and hybridization (Aizen & Rovere, 2010; Bell, Karron, & Mitchell, 2005; Borchsenius, 2002; Campbell, 1985; Morales & Traveset, 2008). Alternatively, flowering times of co-occurring species may *converge* due to facilitative interactions or environmental constraints (Fig. 1b). In this case, the presence of other plant species may increase reproductive success via increased pollinator visitation to collectively larger or more diverse floral displays (Ghazoul, 2006; Gurung, Ratnam, & Ramakrishnan, 2018; Johnson, Peter, Nilsson, & Ågren, 2003; Lopezaraiza–Mikel, Hayes, Whalley, & Memmott, 2007; Moeller, 2004). Synchronous flowering may also decrease the chance of predation on a given species’ flowers and seeds by more broadly spreading the risk across the community (B Rathcke & Lacey, 1985; Beverly Rathcke, 1983). Moreover, phenological character displacement, whether convergent or divergent, is hypothesized to be more likely among closely related species, as more recent ancestry and shared floral morphology make it increasingly likely for taxa to share and experience similar selective pressures from pollinators and predators or experience hybridization and gene flow (W. L. Brown & Wilson, 1956; Darwin, 1859; Levin & Anderson, 1970; Pleasants, 1980; Primack, 1985).

**Figure 1.**
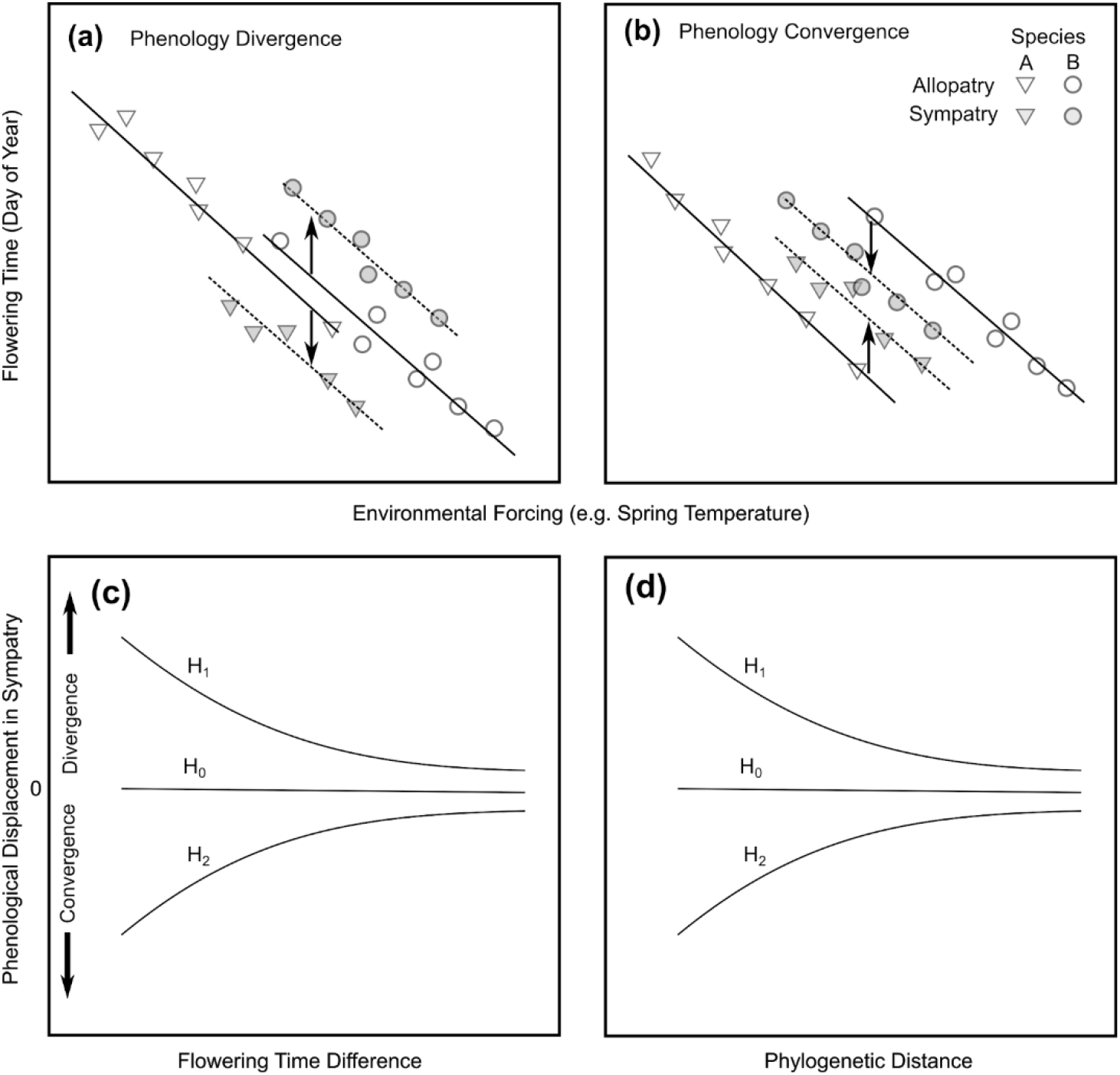
Conceptual framework for assessing phenological displacement. (convergence *or* divergence). Interactions between closely related species can cause phenological traits, here flowering time, to differ between related species growing in sympatry versus those growing in allopatry. For example, if interactions between closely related, co-flowering species are shaped by competition for pollinators or reproductive interference, they may undergo reproductive character divergence in flowering time, causing flowering times to diverge in sympatry (dotted lines) relative to expectations derived from climate-phenology relationships in allopatry (solid lines) (**a**). Alternatively, if interactions are characterized by facilitation or hybridization between species-pairs, then flowering times may converge and be closer in sympatry than in allopatry (**b**). Panels (**c**) and (**d**) show expected patterns across closely related species-pairs under the null hypothesis of no displacement (H0), character divergence (H1), or character convergence (H2). Both H1 and H2 predict larger deviations for sympatric species-pairs that flower at similar times (**c**) and species-pairs that diverged more recently (**d**).

Here, we examine flowering (a)synchrony and evaluate evidence for phenological character displacement across 110 species in 21 diverse families representing major branches of the angiosperm tree of life. We focus on primarily animal-pollinated species, which have been suggested to have more diverse flowering phenologies than wind- or water-pollinated plants (Bolmgren, Eriksson, & Linder, 2003). We used data collected by crowdsourcers from > 42,000 digitized herbarium specimens collected over 120 years and 20 degrees of latitude. We further used these data to examine how flowering phenology has changed over time, and to predict how flowering (a)synchrony among closely related taxa may shift with predicted climatic change.

## Results

Our analysis of 42,805 herbarium specimens collected in the eastern United States from 1881 to 2017 showed substantial variability in mean flowering times and phenological responses to climate, both within and between genera, for our 110 focal species (Fig. 2). Mean flowering dates in climatic conditions typical of the late 20th to early 21st century (1987–2017) varied between 85 DOY and 270 DOY, with a standard deviation of 20 days across species. Using a hierarchical Bayesian linear model (see Methods) we estimated that the mean flowering date of most species (106 of 110) were responsive to spring (March–May) average air temperatures with greater than 90% posterior probability: species flowered an average of 2.5 ± 1.61 (SD) days earlier for every degree of temperature increase. Some species (16 of 110) were also sensitive to spring precipitation, but the average response across all species did not differ from zero (1.7 ± 4.00 days/100 mm of spring precipitation). We found some evidence of phylogenetic signal in peak flowering time (Pagel’s λ = 0.80; p < 0.001) and its sensitivity to spring temperature (Pagel’s λ = 0.54; p < 0.05), but not precipitation (Pagel’s λ = 0; p = 1; Fig. S1). After accounting for temperature and precipitation, a subset of species (18 of 110) also showed credible residual trends over time (i.e., after accounting for shifts in spring temperature or precipitation), flowering on average 0.23 days earlier per decade across all species. Adding additional climatic variables such as summer temperature or vapor pressure deficit failed to improve the overall performance of the model.

**Figure 2.**
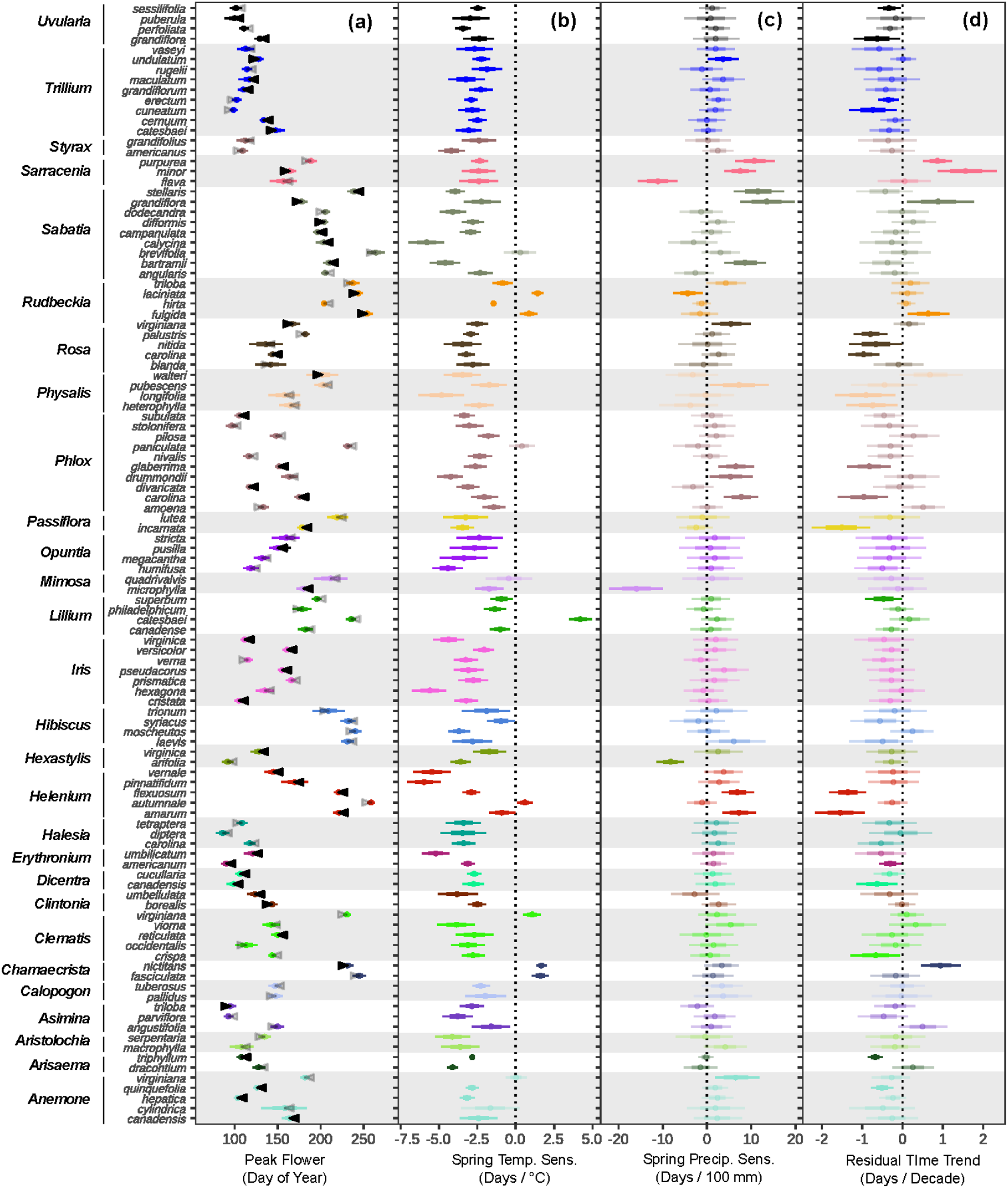
Phenological response summary of 110 angiosperm species. The first column, (**a**), shows estimated mean flowering dates of species spanning 28 genera and 20 plant families during recent climatic conditions (1987-2017), derived from a hierarchical Bayesian linear mixed model. Black arrows indicate significant directional shifts (posterior probability > 90%) in flowering time between 1977 and 2017. Columns (**b**)-(**d**) show estimated climatic sensitivities and residual time trends from the best performing Bayesian hierarchical model of the effects of climate on flowering time. Thick and thin bars represent 50 and 80% credible intervals on the estimates.

Predictions from our hierarchical model additionally allowed us to examine differences in mean flowering dates and assess flowering time convergence or divergence between 74 congener pairs growing in sympatry. On average, species pairs in 24 of 26 genera were not phenologically divergent or convergent relative to null expectations derived from overall climate-phenology relationships (Fig. 3). This was also true overall, with the observed median difference in flowering time across all congener pairs (25 days) virtually identical to the null expectation (24.2 days, Fig. 3 inset). In general, there was no credible phylogenetic signal in patterns of median phenological convergence or divergence between genera, suggesting that patterns of displacement in flowering phenology were not obviously subject to strong evolutionary constraints (Table S1).

**Figure 3.**
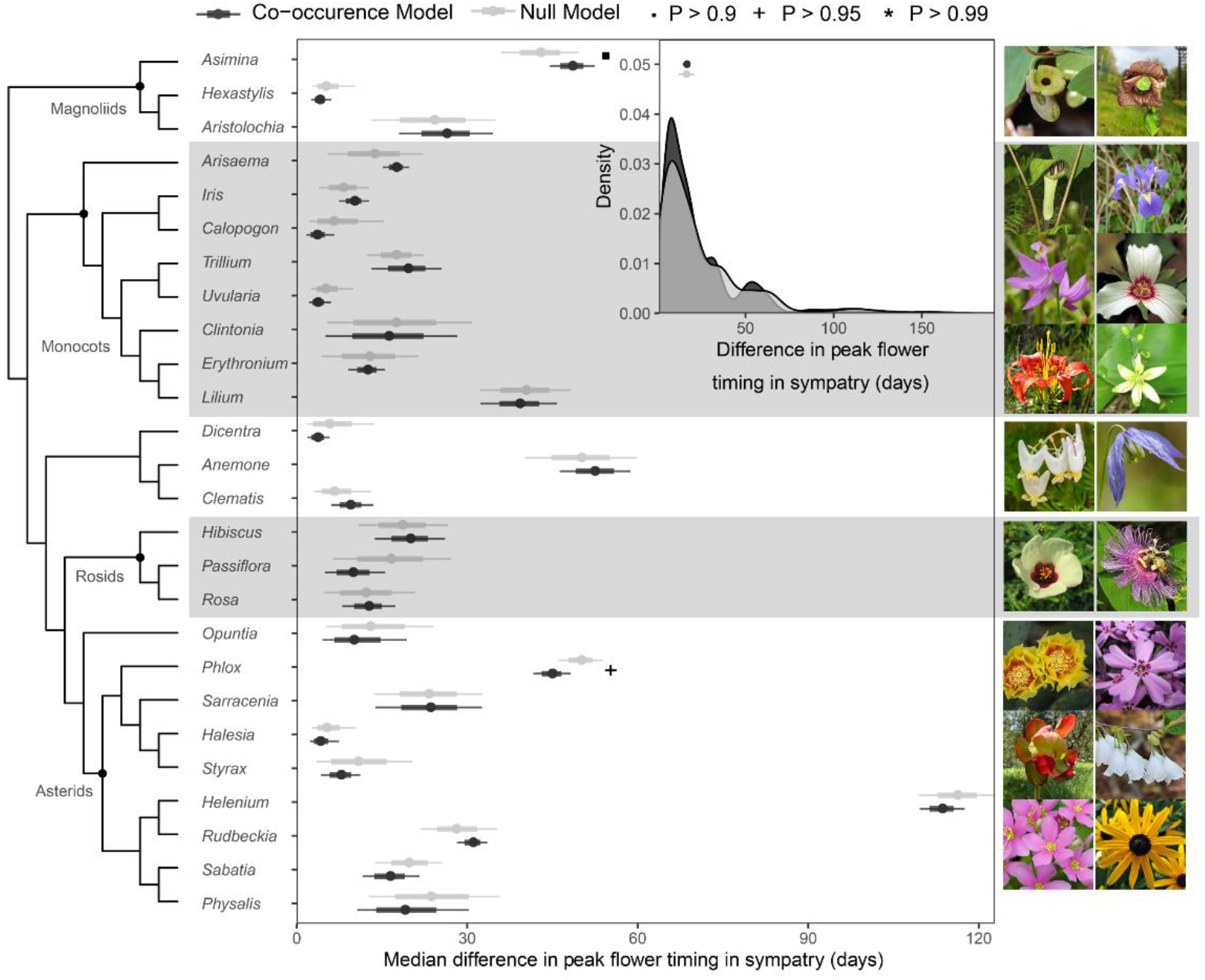
Phenological displacement across genera. Median differences in estimated peak flowering time in sympatry between congener pairs (dark grey) in 28 genera are compared to null expectations that remove the potential influence of species co-occurrence on flowering time (light grey). Density plots (inset) show the distribution of estimates across all congener pairs. Circles and lines at the top left represent estimates and 95% credible intervals, respectively, for the median absolute difference in flowering time across all congener pairs. Genera with median estimates for convergence or divergence that are credibly different from zero are indicated with symbols (· Pr(x ≠ 0) > 0.9; + Pr(x ≠ 0) > 0.95; * Pr(x ≠ 0>0.99)). Major clades are labelled on the phylogeny with black dots. Photographs depicting representative species from each clade are shown to the right. Photographs are from Wikimedia Commons (https://commons.wikimedia.org/) and under a Creative Commons 2.0 generic license. Estimates are derived from a hierarchical Bayesian linear model of flowering time (see Methods).

Most individual co-occurring species pairs did not show large degrees of phenological displacement (Fig. 4). However, we identified highly credible log-linear relationships between the difference in peak flowering time of species pairs and the degree of estimated phenological displacement in sympatry. In terms of days, species pairs with larger differences in the timing of peak flowering displayed greater displacement in their sympatric ranges than those that flower closer in time (Fig. 4a). In contrast, when the degree of phenological displacement was calculated as a percentage of the estimated gap in flowering time, opposite trends emerged (Fig. 4b). Species pairs that tended to flower closer in time displayed greater degrees of displacement in their sympatric ranges relative to the expected gap in their flowering times (Fig. 4b). On average, peak flowering times for species pairs that exhibited phenological convergence were estimated to shift closer by 4.7 ± 0.07 days (22.5 ± 0.67%); pairs that exhibited phenological divergence were estimated to shift 6.1 ± 0.06 days (24.7 ± 0.68%) apart.

**Figure 4.**
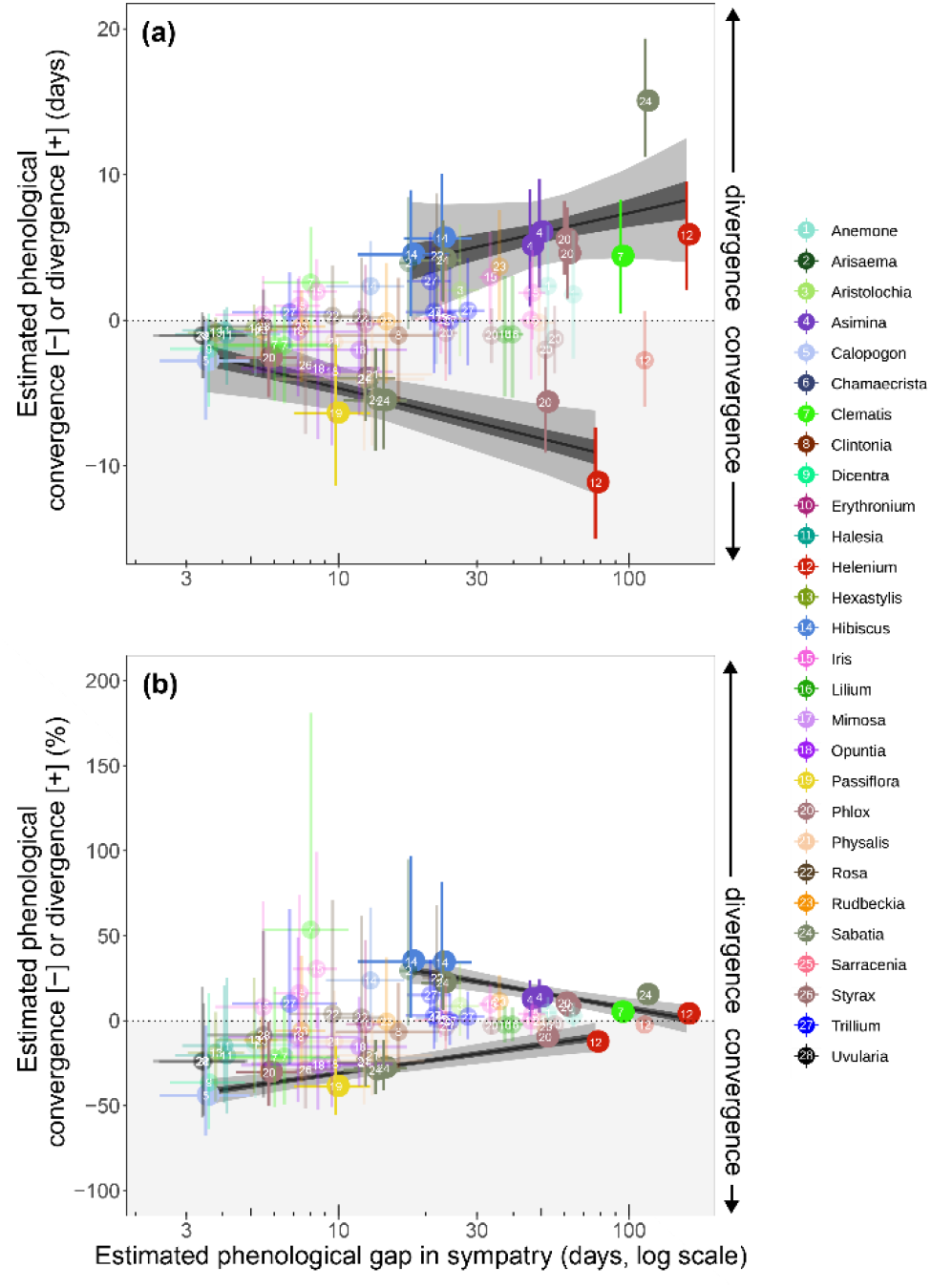
Phenological displacement in sympatry compared to differences in peak flowering time between congener pairs. Estimates of phenological displacement (*y*-axis) are differences in flowering time in sympatry compared to null expectations of flowering time assuming no species interactions, and are depicted as days (**a**) or percent change relative to expected gaps in flowering time among congeners (**b**). Genera appear in different colors and are numbered alphabetically. Circles represent median estimates, and bars represent 25% and 75% posterior quantiles for each species pair. Dark and light shading represents 50% and 95% credible intervals, respectively, for the linear relationships indicated by the black lines.

Since character displacement may be expected to be strongest between more closely related (and thus possibly more ecologically similar) species, we also compared patterns of phenological displacement to phylogenetic distances in a subset of pairs for which we had phylogenetic information. We did not find a credible relationship between pairwise phylogenetic distance and phenological displacement, nor for gaps in peak flowering time (Fig. S2).

To examine how these temporal patterns could change in the near future, we compared the expected timing of peak flowering under climatic conditions of the late 20th century to those expected in the mid-21st century. The flowering season, as defined by the number of days between when 10% and 90% of species pass their peak flower date, was predicted to increase with climatic change by the mid-21st century (Figs. 5a, b). This coincided with an overall expected increase in the temporal gap between peak flowering dates of congeneric species currently growing in sympatry (Figs. 5c, d). For instance, congeneric species in New England and the Atlantic Coastal Plain were projected to flower 2-4 days further apart, on average. In particular, several sympatric species pairs that exhibited convergence in peak flowering time were predicted to experience increased temporal separation in the face of future climate change (Fig. 6).

**Figure 5.**
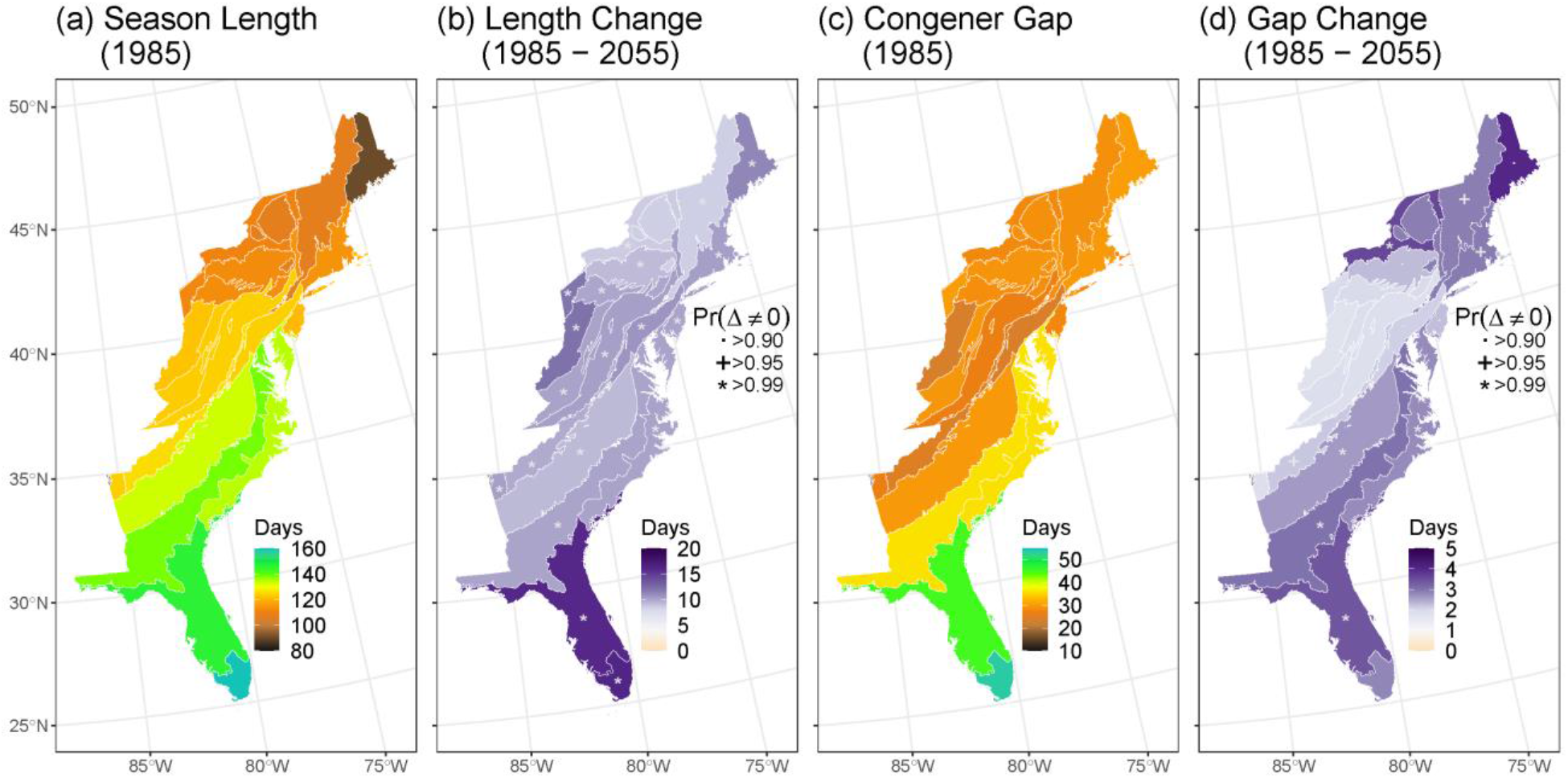
Predicted changes in flowering gaps and season length. Variation in climate-phenology relationships between species and assemblages give rise to large-scale geographic gradients in flowering season length (**a**) and predicted expansion of the flowering season under anthropogenic climate change (**b**). Similar patterns appear in median differences in flowering time between sympatric congeneric pairs (**c**), which are predicted to diverge from each other across much of New England, the Southeastern Coastal Plain, and Peninsular Florida by the mid-21st century (**d**). Maps show county-level predictions from a Bayesian linear mixed model of flowering time summarized by EPA Level III ecoregions (see Methods). Posterior probabilities of changes in growing season length and flowering time for ecoregions in maps (**b**) and (**d**), represented by symbols in each region, are derived from summarizing posterior samples of the Bayesian model.

**Figure 6.**
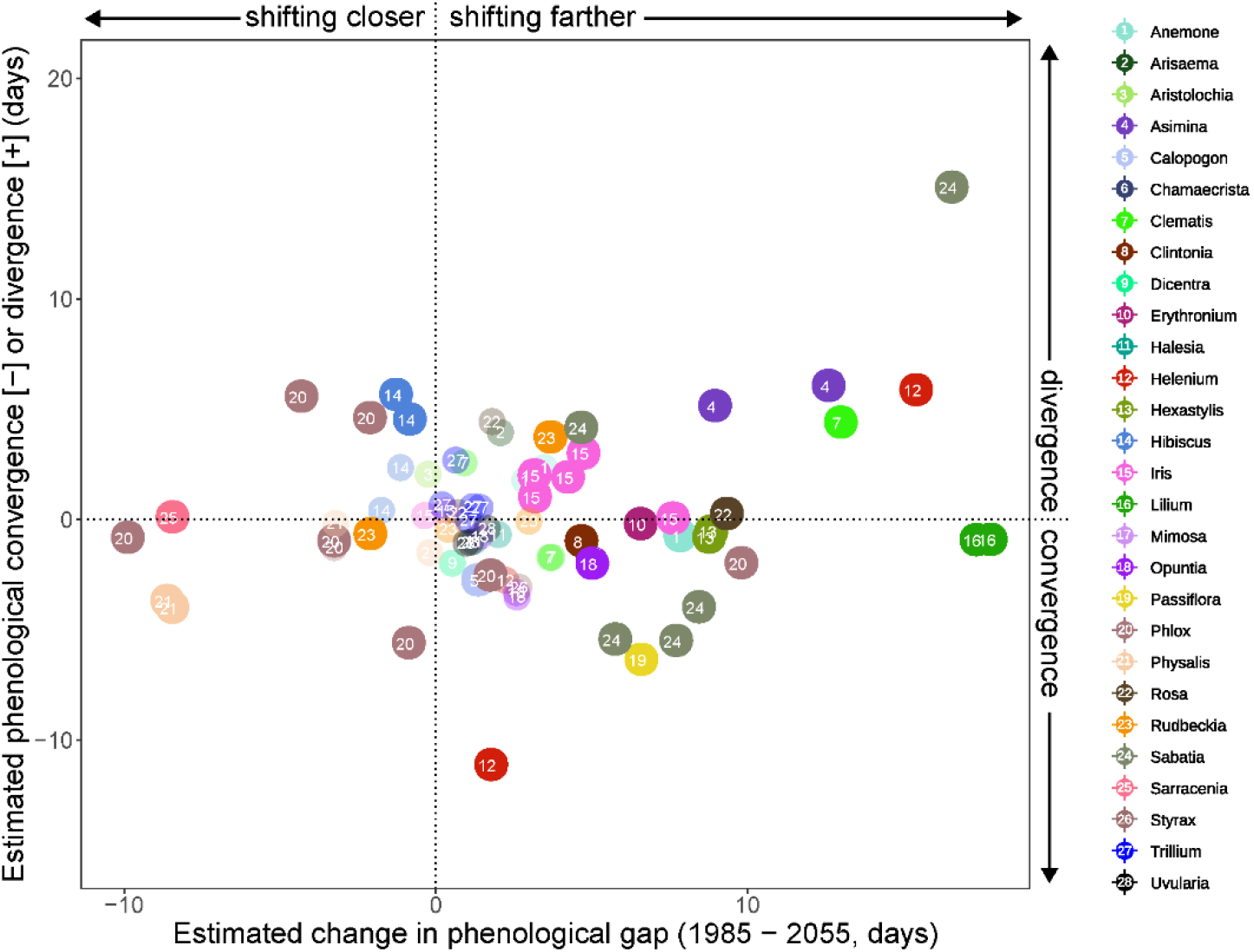
Comparison of expected mid-21st century shifts in flowering synchrony between congener pairs to their degree of phenological displacement in sympatry. Pairs without credible phenological displacement or changes in phenological gaps in synchrony (with lower than 50% posterior probability) are faded. As in Figure 4, colors and numbers within circles represent different genera.

## Discussion

Patterns of flowering time across the landscape result from the dynamic ecological and evolutionary interplay between the phenology of individual taxa and the biotic and abiotic milieu in which they persist (Ackerly, 2003). It has been hypothesized that phenological patterns contributing to the synchronization of reproductive activity with the availability of (a)biotic resources are adaptive (Bolmgren et al., 2003; Brody, 1997; Elzinga et al., 2007) and may be phylogenetically conserved (Kochmer & Handel, 1986). Along these lines, the peak flowering phenology of nearly all the species we examined were sensitive to spring temperatures and we found evidence of phylogenetic signal in both flowering time and its sensitivity to temperature. However, patterns of phenological displacement among closely related, co-occurring species were complex.

### Flowering time displacement in sympatry is not common

On the one hand, sympatric plant species that share pollinators and flower concurrently may reduce each other’s fitness if reproductive success is limited by pollination (Levin & Anderson, 1970; Robertson, 1895). Further, overlapping flowering times between closely related species can result in wasted mating effort or hybrids of reduced fitness (J A Coyne & Orr, 2004). Either of these processes should select for the evolution of staggered, minimally overlapping flowering schedules, especially between closely related taxa. Indeed, divergence in flowering time among sympatric plants have been documented in numerous studies (e.g., (Levin, 2006; Lowry et al., 2008; Spriggs et al., 2019; Stiles, 1977; Stinson, 2004)). On the other hand, phenological convergence can occur if the presence of one species facilitates the reproductive success of another species, or if (a)biotic resources are more temporally constrained in sympatry (Ghazoul, 2006; Beverly Rathcke, 1983).

In contrast to either of these expectations, estimated differences in flowering time varied little for most of the co-occurring congeneric species pairs we examined, regardless of whether they were growing in sympatry or allopatry (Figs. 3, 4). Other taxon-specific studies have also demonstrated a lack of flowering time displacement (usually divergence) at smaller spatial scales (Boulter, Kitching, & Howlett, 2006; Murray et al., 1987). This lack of observed displacement could be the result of at least four factors. First, many congeneric species pairs we examined were effectively isolated in time from each other in terms of peak flowering across their ranges regardless of co-occurrence (Fig. 2a). In such cases, small shifts in phenology likely would have negligible effects on competitive or facilitative interactions among co-occurring taxa. Second, in many systems pollinators are not as limiting as other essential resources (Horvitz & Schemske, 1988; B Rathcke & Lacey, 1985). Third, the direction, intensity, and outcome of reproductive interactions may vary at smaller spatial scales, mitigated by the abundance and density of interacting species, none of which our large-scale analyses could detect. Fourth, and finally, flowering-time displacement is but one of several mechanisms that can either reduce interspecific competition and gene flow or facilitate net reproductive gains (Elzinga et al., 2007; Levin, 1971; Moeller, 2004).

### Phenological displacement is greater among species that flower close in time

Among species pairs for which we did observe phenological displacement, there was a highly credible log-linear relationship between the difference in peak flowering time of species pairs and the degree of estimated phenological displacement in sympatry. Congener pairs that flowered further apart in time displayed greater degrees of displacement in terms of number of days converged or diverged in sympatry. This result may reflect that opportunities for possible adaptive or stochastic shifts is associated with increases in time between flowering events. Further, as species flowering further apart are less likely to interact directly, phenological shifts may have little effect on reproductive competition or facilitation. For instance, a convergence of 10 days for a pair of *Helenium* species that tend to flower apart by 3 months is unlikely to greatly alter the nature of their interactions. Thus, it is more informative to examine the degree of displacement in the context of overall flowering time difference.

When we quantified the relationship between proportional phenological displacement to estimated gaps in flowering time, however, we found that the amount of displacement was greater among species pairs that tended to flower closer in time. In particular, closely related species with similar peak flowering times tended to exhibit even more convergent flowering times when they co-occurred. This observation supports hypotheses that aggregated flowering of species during a relatively narrow window of time can be advantageous in certain conditions (B Rathcke & Lacey, 1985; Thomson, 1978).

However, flowering phenology can be influenced by other selective pressures as well. For instance, selection to avoid herbivores can conflict with pollinator-mediated selection on flowering time (Elzinga et al., 2007; Sletvold et al., 2015). In certain regions, climatic conditions suitable for flowering may be short-lived, resulting in phenological convergence among lineages (Levin, 2006). Edaphic conditions also can mediate phenological responses (Brady, Kruckeberg, & Bradshaw Jr, 2005; Sambatti & Rice, 2007). Flowering time also can be constrained indirectly by selection effects on the timing of germination or dispersal (Primack, 1987).

In summary, our results suggest that while the direction and degree of displacement varies greatly among taxa, displacement is typically stronger among species that flower closer in time. However, we did not detect any relationship between displacement and phylogenetic distance, suggesting that the strength of inter-specific interactions do not scale predictably with evolutionary relatedness. Relatedness is not always a good predictor of the strength of inter-specific interactions (Bennett, Lamb, Hall, Cardinal-McTeague, & Cahill, 2013; Cahill Jr, Kembel, Lamb, & Keddy, 2008), but future studies incorporating a more comprehensive phylogenetic framework are necessary to elucidate whether this is indeed the case for phenological displacement (Davis, Willis, Primack, & Miller-Rushing, 2010).

### Climate change will alter temporal interactions among closely related species

As the climate continues to change, the diverse competitive or facilitative outcomes among species will be driven in part by idiosyncratic shifts in phenology. For instance, if the lack of flowering-time divergence among closely related sympatric species is at least partially the result of facilitative interactions among taxa, there may be negative consequences of future divergence. Less diverse, smaller floral displays may reduce pollinator visitation, whereas increased asynchrony in flowering can concentrate the chance of predation on a given species’ reproductive organs (Feldman, Morris, & Wilson, 2004; Ghazoul, 2006; Gurung et al., 2018; Moeller, 2004; Beverly Rathcke, 1983). Phenological divergence also can create new reproductive niches, which may be conducive to invasion by non-native species (Sherry et al., 2007; Wolkovich & Cleland, 2014). Finally, changes in climate can directly modify selective pressures on flowering phenology and alter associated biotic interactions across trophic levels (Filchak, Roethele, & Feder, 2000; Forkner, Marquis, Lill, & CORFF, 2008; Franks et al., 2007; Renner & Zohner, 2018). Although it is difficult to predict the outcome of increased divergence of flowering times between co-occurring closely related, species, climate-induced changes in phenology will lead to new temporal patterns of reproductive overlap, potentially affecting species interactions, and result in altered species compositions across space and time (Franks & Weis, 2009; Pau et al., 2011; Post, Forchhammer, Stenseth, & Callaghan, 2001; Sherry et al., 2007; Waser & Real, 1979).

Recognizing that some cases of true phenological character displacement do exist, future assessments should seek to understand how flowering time interacts with other ecological and evolutionary constraints such as pollinator availability and postzygotic reproductive barriers. Although our study focused on temperate, insect-pollinated plants, we included a wide array of species from across the angiosperm phylogeny, ranging from trees to understory herbs. The same methods could be used to test whether similar patterns are found for wind-pollinated plants, among which it has been suggested that flowering time displacement could be more common (Hopkins, 2013; McNeilly & Antonovics, 1968; Savolainen et al., 2006). The methods and results presented here provides one promising path towards understanding how the phenological landscape is structured and may respond to future environmental change.

## Materials and Methods

### Selection of species and specimens

We used digitized specimens from two of the most comprehensive digitized regional floras in the world, the Consortium of Northeastern Herbaria (CNH; http://portal.neherbaria.org/portal/) and Southeast Regional Network of Expertise and Collections (SERNEC; http://sernecportal.org/portal/index.php). We selected animal-pollinated species from across the eastern United States that satisfied the following criteria: (i) included collection dates and at least county-level locality data; (ii) comprised at least 50 unique collections across space and time; (iii) had reproductive structures (i.e., buds, flowers, and fruit) that were easily identifiable and quantifiable by crowdsourcers; and (iv) had at least one other congeneric species with a partially overlapping geographic range in our study area. Citizen-scientists hired through Amazon’s Mechanical Turk service (MTurk; https://www.mturk.com/) counted the number of buds, flowers, and fruits to assess peak flowering time. See Park *et al*. (Park et al., 2018) for detailed crowdsourcing methods. Our final dataset comprised 110 species in 28 genera across 21 angiosperm families (Table S2). As our specimen data alone gave an incomplete picture of species county-level distributions, we determined co-occurrence among congener groups based on combining county-level distributions from our specimen data with county checklist data from the United States Department of Agriculture PLANTS Database (https://plants.usda.gov/).

We used estimates of historic (1895–2017) average monthly air temperature and precipitation data at 2.5 arc-minute resolution from PRISM (product AN81m; http://prism.oregonstate.edu/). Accurate locality data were not available for the majority of historic specimen records (Park & Davis, 2017), so we used county as our geographical unit of analysis. For each county and year, we estimated the mean monthly temperature, precipitation and elevation, and assigned these values to each specimen. Though counties can vary in size and climate, counties in states along the east coast of the United States are generally small in size and geographically homogeneous, and within-county variation in climate does not significantly affect estimations of phenological response in this area (Park et al., 2018).

### Data collection

Crowdsourcers used the citizen science platform *CrowdCurio* (Willis et al., 2017) to gather phenological information from over 40,000 digitized herbarium specimens collected over 120 years and 20 degrees of latitude. The expansive spatial, temporal, and phylogenetic sampling offered by herbarium collections has become increasingly accessible with widespread digitization (Hedrick et al., 2020) and crowdsourcing has been demonstrated to be an effective, reliable method for assessing phenological traits from natural history collections (Willis et al., 2017). The flowering patterns derived from specimens have been shown to reflect those assessed from field surveys (Borchert, Meyer, Felger, & Porter-Bolland, 2004; Davis et al., 2015). Further, specimens allow us to assess phenological community patterns at macroecological scales essential to obtain a generalizable understanding of the phenological responses of species and communities (Doi, Gordo, Mori, & Kubo, 2017). From the *CrowdCurio-derived* observations, we first computed the median number of buds, flowers, and fruits observed on each herbarium specimen. For phenological analysis, we used specimens that met the following criteria: (i) contained at least one open flower, (ii) contained more flowers than the combined number of buds and fruits, (iii) contained a number of flowers representing at least 5% of the maximum (95th quantile) number of flowers observed on a given species, and (iv) had collection dates ≥ the 5th quantile and ≤ the 95th quantile of flowering dates. These filters ensured that the specimens used for analysis were in full flower and excluded outlier specimens collected outside of the main flowering period of each species. Of the 42,805 specimens that were originally phenotyped, we used 19,543 in our hierarchical model of flowering time. Although our filtering strategy was quite aggressive, we verified that including less aggressive filters (i.e., removing filters ii-iv above) did not qualitatively alter our results.

### Statistical Modeling

Bayesian hierarchical models can help to overcome common biases inherent with herbarium data (Park et al., 2018). For example, specimen data are spatiotemporally sparse, phenological traits are highly plastic, and estimates of displacement among species pairs within a given clade are not independent of one another (Daru et al., 2018; Theobald, Breckheimer, & HilleRisLambers, 2017). Relatively few specimens in our dataset were collected at the same locality and in the same year as their congeners. Flowering time for many of our focal species is highly sensitive to environmental forcing (warmer spring temperatures generally inducing earlier flowering) and flowering times sometimes differed across species’ ranges because of climatic differences unrelated to interspecific interactions.

#### Model overview

Our Bayesian model first involved applying a single hierarchical linear model to the filtered specimen dataset to predict the mean flowering date of each species from climate and co-occurring congeners. We then used posterior samples from this model to generate predictions of flowering time with and without terms representing the influence of congeneric species on flowering time. These predictions allowed us to estimate differences in mean flowering time in sympatry for each species pair that were associated with the presence or absence of a particular congener, and separate them from differences in flowering time resulting from underlying species-specific relationships between phenology and climate. Generating estimates from each posterior sample of the model allowed us to propagate uncertainty in estimates of species-specific climate and congener effects to our pairwise estimates of phenological divergence and overall estimates across all species pairs and relationships between divergence, mean flowering time, and phylogenetic distance.

#### Statistical model of flowering time

To estimate species-specific flowering times, and the effects of climate and congeners on the phenology of each focal species, we fit a hierarchical Bayesian linear regression model. The model treated the day of year (DOY) recorded on each flowering specimen as a normally distributed random variable with mean μ_*DOY*_ and standard deviation σ_*DOY*_. Mean flowering date was related to spring (March - May) average air temperature (*T*) in the county (*c*) and year (*y*) that the specimen was collected using a linear function with species-specific intercepts (β0_*j*_) and slopes (β1_*j*_). The model also includes separate categorical intercept terms for each county (β2_*c*_), genus (β3_*g*_), and congener group (β4_*u*_):

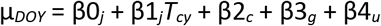

All beta parameters were drawn from normal distributions with hyperparameters:

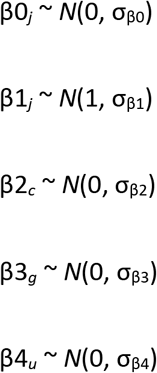

Air temperature estimates were derived from the PRISM 2.5 arc-minute gridded data as listed above. Terms for genus (3g) were included to account for the potential non-independence of phenology within genera. The congener group variable (*g*), was a categorical variable with unique values indexing different combinations of congeneric species that occur in different parts of a species range. For example, if species A co-occurred with only congener B in county 1 and county 2 but congeners B and C in county 3, then the indices for this variable would be 1, 1, and 2, respectively. The estimates of these coefficients should capture the combined influence of co-occurring congeneric species on flowering time. We fit our model using Hamiltonian Monte Carlo sampling techniques implemented using the *rstanarm* package ver. 2.19.3 (Goodrich, Gabry, Ali, & Brilleman, 2020) in R version 3.6.3 (R Core Team, 2020). The model was fit using four sampling chains of 8000 iterations each, with the last 1000 iterations retained. We verified model convergence and desirable sampler behavior by visually assessing the model fit using functions implemented in the *bayesplot* package ver. 1.7.1 (Gabry & Mahr, 2019), as well as the Gelman-Rubin statistic (Gelman & Rubin, 1992). The effective sample size for all parameters was greater than 1000. To assess model fit and ensure that samples from the posterior predictive distribution of the model closely resembled the real data, we used the built-in predictive checks in *rstanarm*.

#### Estimating flowering displacement in sympatry

We processed posterior samples from our model to generate estimates of differences in mean flowering time in sympatry across all congeneric species pairs where we had observations of co-occurrence (sympatry) and non-co-occurrence (allopatry) across at least 3 different counties each. For each sympatric congener pair, we used the complete fit model described above to generate estimates of flowering time for each focal species and each congener in each county and year where we had specimens of the focal species and we had either specimens or checklist records of the presence of its congener (in any year). These estimates of flowering time in sympatry (co-occurrence estimates) incorporate model terms representing species-specific flowering times (β0_*j*_, β3_*g*_), and climate-phenology relationships (β1_*j*_), and, critically, the effects of co-occurring congeners (β4_*u*_). We then subtracted the predicted flowering times for focal species from flowering time estimates of their congener pair and took the absolute value to generate an estimate of the difference in flowering time for each congener pair in each sympatric county in each year where we had specimens of the focal species. Finally, to represent a typical difference in flowering time in sympatry, we computed the median difference in flowering time across all sympatric counties for each congener pair. To estimate uncertainty in flowering times in our co-flowering estimates, we generated 4000 estimates of each pairwise median, one from each posterior sample of our model.

To isolate the influence of sympatry itself on differences in flowering time, we also generated flowering time estimates for congener pairs that exclude terms representing the influence of co-occurrence, estimates (null expectation). To accomplish this, we used an approach identical to the one we describe above (to generate the co-occurrence estimates) with one key difference: predictions did not include the model’s co-occurrence terms (β4_*u*_) for either species. Subtracting differences in flowering time of the co-occurrence estimate from the null estimate allows us to measure how much co-occurrence with congeners might affect differences in flowering time, which we define as phenological displacement in sympatry. This was done for each iteration of our Bayesian model, which properly propagated uncertainty from the original data to our final estimates of phenological displacement, both for overall estimates across all species pairs at the genus level (Fig. 3) and individual pairwise comparisons (Fig. 4).

#### Testing predictions of phenological character displacement

Our two alternative hypotheses, that reproductive interference and pollinator competition drive phenological character displacement (Fig. 1, H1) or that facilitative interactions or environmental constraints drive phenological convergence (Fig. 1, H2) make several testable predictions regarding patterns of co-flowering among species pairs. Both hypotheses lead to the prediction that gaps in flowering time of species pairs in sympatry will differ from expectations derived from underlying climate-phenology relationships (i.e., divergences in sympatry credibly different from zero), and these deviations will be larger for species pairs that flower at similar times and species pairs that are closely related. We tested these predictions by comparing our estimates of phenological displacement in sympatry, to differences in mean flowering time in a hypothetical common-garden setting, and phylogenetic distances. Phylogenetic distances were calculated from a set of published time-calibrated phylogenies of the North American flora based on twelve commonly used molecular loci (Park et al., 2020). Of the 110 species examined, 85 were represented on the phylogeny, and we were able to calculate phylogenetic distance between 48 of the 74 co-occurring congener pairs. Differences in mean flowering time for each species pair were taken from the null estimates described above. For each of 1000 posterior samples of our model, we recorded how many showed a negative slope in the linear relationship between (log-transformed) flowering time differences and estimates of phenological displacement in sympatry across all species pairs. Although we did not have posterior samples for phylogenetic distances, we used 100 dated bootstrap replicates in a similar fashion, comparing them to posterior samples of phenological displacement and recording how many posterior samples out of 1000 showed the expected negative relationship between phenological displacement and phylogenetic distance.

To examine how gaps in peak flowering time will shift with climatic change in the near future, we compared the expected timing of peak flowering under climatic conditions of the late 20th century to those expected in the mid-21st century. Predictions for 1985 used mean environmental conditions (1970–1999 spring temperature and precipitation) as estimated from PRISM. Mid-21st century (2055) predictions used county-level temperature and precipitation change estimates (2040–2069) from a set of 18 Coupled Model Inter-comparison Project 5 (CMIP5) global circulation models downscaled and summarized to the county level using the Multivariate Adaptive Constructed Analogs (MACA) algorithm (Elias et al., 2018). Although these predictions are for a high-emissions scenario (RCP 8.5), predictions for different emissions scenarios do not diverge substantially until the late-21st century.

## Data availability

All data, (permanent links to) imagery, and model code are available from the Harvard Forest Data Archive (https://harvardforest.fas.harvard.edu/data-archive), number HF335 (doi:10.6073/pasta/c17fcc2ba0f9212938b2b5f6161615d8), and from the Environmental Data Initiative (will provide DOI following accession).

## Acknowledgements

We express gratitude to the many collectors and curators of biodiversity data who made this research possible. We also thank members of the Plants & Climate discussion group for their invaluable insights and comments. This study was funded as part of the New England Vascular Plant Project to CCD (NSF-DBI: EF1208835), NSF-DEB 1754584 to CCD, DSP, and AME. AME’s participation in this project was supported by Harvard Forest.

## Author Contributions

DSP and CCD conceived the study; DSP, CCD, AME, and IKB designed the study; DSP and GML collected data; IKB, AME, and DSP analyzed the data; DSP and IBK drafted the first version of the manuscript and all authors contributed significantly to subsequent revisions.

## Supplementary Information

**Figure S1.** Phylogenetic signal in peak flowering phenology and its sensitivity to environmental forcings.

**Figure S2.** Estimated phenological gap and displacement in sympatry compared to phylogenetic distances between congener pairs.

**Table S1.** Phylogenetic signal in patterns of median phenological displacement between genera.

**Table S2.** List of species used in study.

