## supplemental information for "Phenological displacement is uncommon among sympatric angiosperms"

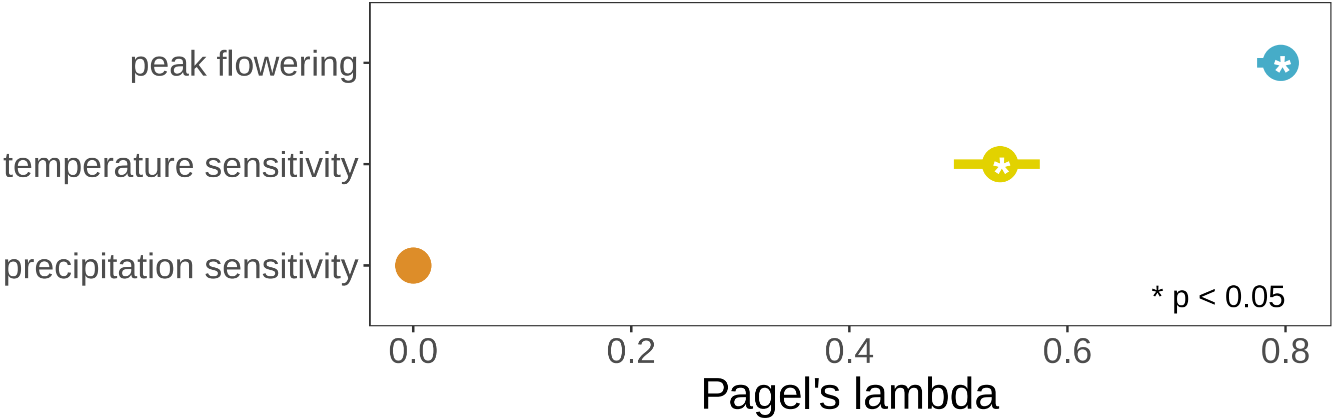


**Figure S1.** Phylogenetic signal in peak flowering phenology and its sensitivity to environmental forcings. Points indicate values of Pagel's lambda assessed on the maximum likelihood phylogeny and bars represent values from 100 bootstrap replicate trees.

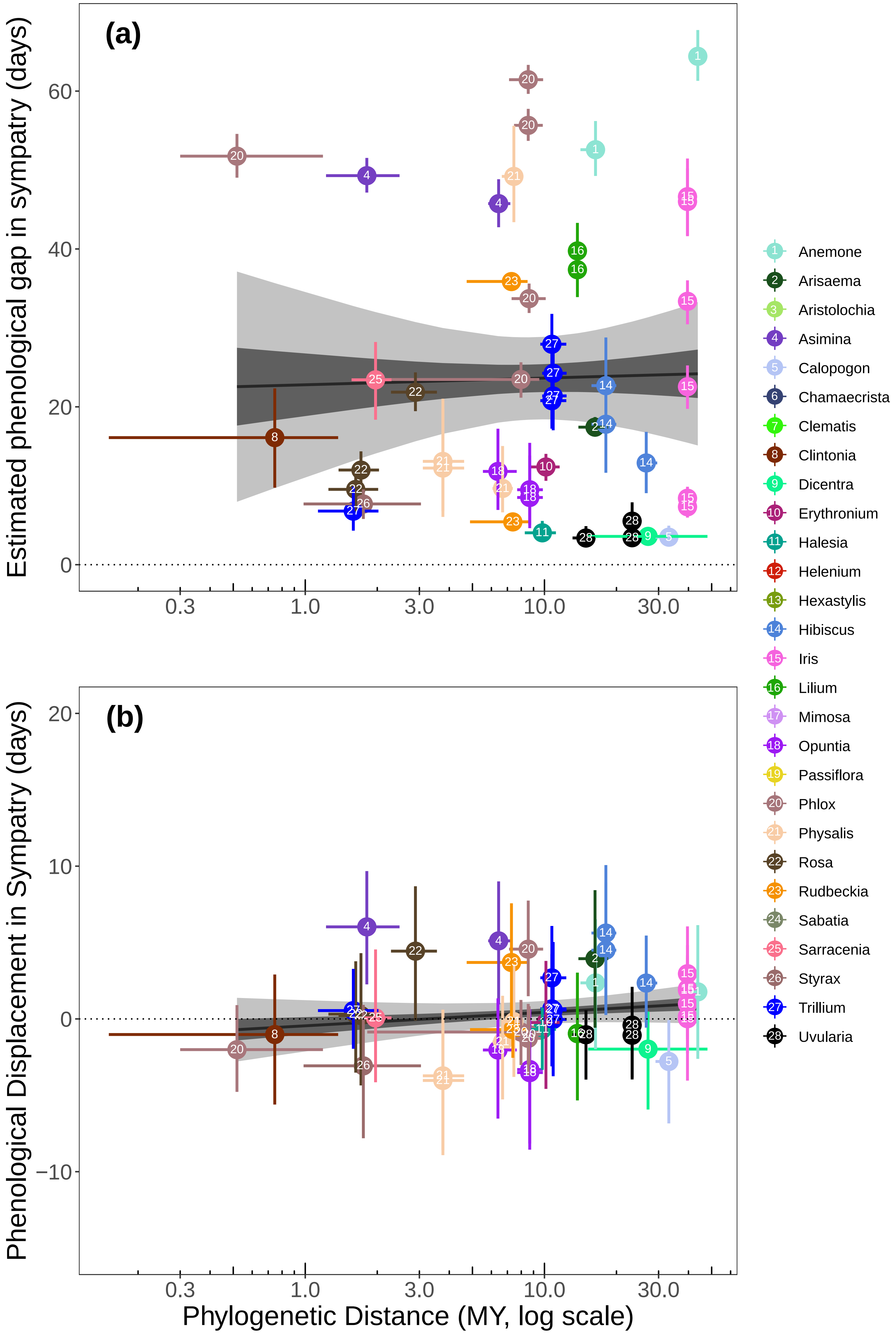


**Figure S2.** Estimated phenological gap (a) and displacement (b) in sympatry compared to phylogenetic distances between congener pairs. Estimates of phenological displacement are differences in flowering time in sympatry compared to null expectations of flowering time assuming no species interactions. Genera appear in different colors and are numbered alphabetically. Circles represent median estimates, and bars represent 25% and 75% posterior quantiles for each species pair. In the case of phylogenetic distances, bars represent 25% and 75% quantiles on distances derived from bootstrap replicates of the underlying phylogenetic tree. Dark and light shading represents 50% and 95% credible intervals, respectively, for the linear relationships indicated by the black lines.

**Table S1.** Phylogenetic signal in patterns of median phenological displacement between genera.

| tree | lambda | p-value |
|---|---|---|
| ML | 0.501888 | 0.302722 |
| bootstrap1 | 0.518605 | 0.35225 |
| bootstrap2 | 0.543509 | 0.305458 |
| bootstrap3 | 0.521176 | 0.303999 |
| bootstrap4 | 0.580856 | 0.279425 |
| bootstrap5 | 0.534919 | 0.284567 |
| bootstrap6 | 0.538135 | 0.286899 |
| bootstrap7 | 0.472953 | 0.302022 |
| bootstrap8 | 0.457321 | 0.330035 |
| bootstrap9 | 0.527478 | 0.29437 |
| bootstrap10 | 0.526716 | 0.288677 |
| bootstrap11 | 0.490593 | 0.314795 |
| bootstrap12 | 0.480321 | 0.312284 |
| bootstrap13 | 0.552671 | 0.294576 |
| bootstrap14 | 0.50509 | 0.289434 |
| bootstrap15 | 0.503175 | 0.292718 |
| bootstrap16 | 0.497726 | 0.395374 |
| bootstrap17 | 0.524045 | 0.372405 |
| bootstrap18 | 0.451695 | 0.305484 |
| bootstrap19 | 0.500506 | 0.297021 |
| bootstrap20 | 0.499593 | 0.364476 |
| bootstrap21 | 0.49107 | 0.304401 |
| bootstrap22 | 0.582059 | 0.258702 |
| bootstrap23 | 0.542523 | 0.368822 |
| bootstrap24 | 0.508945 | 0.341359 |
| bootstrap25 | 0.556489 | 0.285953 |
| bootstrap26 | 0.506415 | 0.300887 |
| bootstrap27 | 0.50539 | 0.365222 |
| bootstrap28 | 0.516526 | 0.301752 |
| bootstrap29 | 0.538637 | 0.290773 |
| bootstrap30 | 0.499252 | 0.305341 |
| bootstrap31 | 0.523236 | 0.367154 |
| bootstrap32 | 0.493602 | 0.3176 |
| bootstrap33 | 0.559693 | 0.267395 |
| bootstrap34 | 0.570152 | 0.30632 |
| bootstrap35 | 0.511541 | 0.30152 |
| bootstrap36 | 0.534109 | 0.28931 |
| bootstrap37 | 0.516972 | 0.376633 |
| bootstrap38 | 0.52911 | 0.307283 |

| | | |
|---|---|---|
| bootstrap39 | 0.460882 | 0.314531 |
| bootstrap40 | 0.501266 | 0.30953 |
| bootstrap41 | 0.519405 | 0.27601 |
| bootstrap42 | 0.54891 | 0.364962 |
| bootstrap43 | 0.52585 | 0.292092 |
| bootstrap44 | 0.50258 | 0.311295 |
| bootstrap45 | 0.502974 | 0.298144 |
| bootstrap46 | 0.515205 | 0.288923 |
| bootstrap47 | 0.514185 | 0.300465 |
| bootstrap48 | 0.475467 | 0.307913 |
| bootstrap49 | 0.484253 | 0.3054 |
| bootstrap50 | 0.533796 | 0.296216 |
| bootstrap51 | 0.513038 | 0.305667 |
| bootstrap52 | 0.530194 | 0.290785 |
| bootstrap53 | 0.562219 | 0.287779 |
| bootstrap54 | 0.500898 | 0.309254 |
| bootstrap55 | 0.526793 | 0.298868 |
| bootstrap56 | 0.530021 | 0.290039 |
| bootstrap57 | 0.524131 | 0.311092 |
| bootstrap58 | 0.525664 | 0.291347 |
| bootstrap59 | 0.518521 | 0.281559 |
| bootstrap60 | 0.521619 | 0.299744 |
| bootstrap61 | 0.543227 | 0.296341 |
| bootstrap62 | 0.533235 | 0.303972 |
| bootstrap63 | 0.49543 | 0.311017 |
| bootstrap64 | 0.525289 | 0.357968 |
| bootstrap65 | 0.545282 | 0.305758 |
| bootstrap66 | 0.544941 | 0.281392 |
| bootstrap67 | 0.480279 | 0.398659 |
| bootstrap68 | 0.53405 | 0.296675 |
| bootstrap69 | 0.521441 | 0.324474 |
| bootstrap70 | 0.493323 | 0.300252 |
| bootstrap71 | 0.511695 | 0.289869 |
| bootstrap72 | 0.558266 | 0.292388 |
| bootstrap73 | 0.47417 | 0.31351 |
| bootstrap74 | 0.507268 | 0.300959 |
| bootstrap75 | 0.521843 | 0.284838 |
| bootstrap76 | 0.494166 | 0.29701 |
| bootstrap77 | 0.54843 | 0.318719 |
| bootstrap78 | 0.48244 | 0.316815 |
| bootstrap79 | 0.521189 | 0.335817 |

| | | |
|---|---|---|
| bootstrap80 | 0.542574 | 0.289319 |
| bootstrap81 | 0.491471 | 0.410231 |
| bootstrap82 | 0.559104 | 0.274712 |
| bootstrap83 | 0.542718 | 0.347602 |
| bootstrap84 | 0.477873 | 0.471148 |
| bootstrap85 | 0.516159 | 0.300204 |
| bootstrap86 | 0.497054 | 0.36825 |
| bootstrap87 | 0.493454 | 0.30163 |
| bootstrap88 | 0.468077 | 0.322953 |
| bootstrap89 | 0.493523 | 0.304244 |
| bootstrap90 | 0.51131 | 0.302388 |
| bootstrap91 | 0.493885 | 0.299716 |
| bootstrap92 | 0.519211 | 0.311696 |
| bootstrap93 | 0.501325 | 0.311841 |
| bootstrap94 | 7.90E-05 | 1 |
| bootstrap95 | 0.529476 | 0.297723 |
| bootstrap96 | 0.490968 | 0.304743 |
| bootstrap97 | 0.506295 | 0.32336 |
| bootstrap98 | 0.519866 | 0.293525 |
| bootstrap99 | 0.482639 | 0.33721 |
| bootstrap100 | 0.506659 | 0.294695 |

**Table S2.** List of species used in study.

| Order | Family | Genus | Species |
|---|---|---|---|
| Asterales | Asteraceae | *Helenium* | *Helenium amarum* |
| Asterales | Asteraceae | *Helenium* | *Helenium autumnale* |
| Asterales | Asteraceae | *Helenium* | *Helenium flexuosum* |
| Asterales | Asteraceae | *Helenium* | *Helenium pinnatifidum* |
| Asterales | Asteraceae | *Helenium* | *Helenium vernale* |
| Asterales | Asteraceae | *Rudbeckia* | *Rudbeckia fulgida* |
| Asterales | Asteraceae | *Rudbeckia* | *Rudbeckia hirta* |
| Asterales | Asteraceae | *Rudbeckia* | *Rudbeckia laciniate* |
| Asterales | Asteraceae | *Rudbeckia* | *Rudbeckia triloba* |
| Ericales | Polemoniaceae | *Phlox* | *Phlox amoena* |
| Ericales | Polemoniaceae | *Phlox* | *Phlox carolina* |
| Ericales | Polemoniaceae | *Phlox* | *Phlox divaricata* |
| Ericales | Polemoniaceae | *Phlox* | *Phlox drummondii* |
| Ericales | Polemoniaceae | *Phlox* | *Phlox glaberrima* |
| Ericales | Polemoniaceae | *Phlox* | *Phlox nivalis* |
| Ericales | Polemoniaceae | *Phlox* | *Phlox paniculata* |
| Ericales | Polemoniaceae | *Phlox* | *Phlox pilosa* |
| Ericales | Polemoniaceae | *Phlox* | *Phlox stolonifera* |
| Ericales | Polemoniaceae | *Phlox* | *Phlox subulata* |
| Ranunculales | Papaveraceae | *Dicentra* | *Dicentra canadensis* |
| Ranunculales | Papaveraceae | *Dicentra* | *Dicentra cucullaria* |
| Ranunculales | Ranunculaceae | *Anemone* | *Anemone canadensis* |
| Ranunculales | Ranunculaceae | *Anemone* | *Anemone cylindrica* |
| Ranunculales | Ranunculaceae | *Anemone* | *Anemone hepatica* |
| Ranunculales | Ranunculaceae | *Anemone* | *Anemone quinquefolia* |
| Ranunculales | Ranunculaceae | *Anemone* | *Anemone virginiana* |
| Ranunculales | Ranunculaceae | *Clematis* | *Clematis crispa* |
| Ranunculales | Ranunculaceae | *Clematis* | *Clematis occidentalis* |
| Ranunculales | Ranunculaceae | *Clematis* | *Clematis reticulata* |
| Ranunculales | Ranunculaceae | *Clematis* | *Clematis viorna* |
| Ranunculales | Ranunculaceae | *Clematis* | *Clematis virginiana* |
| Alismatales | Araceae | *Arisaema* | *Arisaema dracontium* |
| Alismatales | Araceae | *Arisaema* | *Arisaema triphyllum* |
| Asparagales | Orchidaceae | *Calopogon* | *Calopogon pallidus* |
| Asparagales | Orchidaceae | *Calopogon* | *Calopogon tuberosus* |
| Asparagales | Iridaceae | *Iris* | *Iris cristata* |
| Asparagales | Iridaceae | *Iris* | *Iris hexagona* |
| Asparagales | Iridaceae | *Iris* | *Iris prismatica* |
| Asparagales | Iridaceae | *Iris* | *Iris pseudacorus* |
| Asparagales | Iridaceae | *Iris* | *Iris verna* |
| Asparagales | Iridaceae | *Iris* | *Iris versicolor* |
| Asparagales | Iridaceae | *Iris* | *Iris virginica* |
| Caryophyllales | Cactaceae | *Opuntia* | *Opuntia humifusa* |
| Caryophyllales | Cactaceae | *Opuntia* | *Opuntia megacantha* |
| Caryophyllales | Cactaceae | *Opuntia* | *Opuntia pusilla* |
| Caryophyllales | Cactaceae | *Opuntia* | *Opuntia stricta* |
| Ericales | Sarraceniaceae | *Sarracenia* | *Sarracenia flava* |
| Ericales | Sarraceniaceae | *Sarracenia* | *Sarracenia minor* |
| Ericales | Sarraceniaceae | *Sarracenia* | *Sarracenia purpurea* |
| Ericales | Styracaceae | *Halesia* | *Halesia carolina* |
| Ericales | Styracaceae | *Halesia* | *Halesia diptera* |
| Ericales | Styracaceae | *Halesia* | *Halesia tetraptera* |
| Ericales | Styracaceae | *Styrax* | *Styrax americanus* |
| Ericales | Styracaceae | *Styrax* | *Styrax grandifolius* |
| Fabales | Fabaceae | *Chamaecrista* | *Chamaecrista fasciculata* |
| Fabales | Fabaceae | *Chamaecrista* | *Chamaecrista nictitans* |
| Fabales | Fabaceae | *Mimosa* | *Mimosa microphylla* |
| Fabales | Fabaceae | *Mimosa* | *Mimosa quadrivalvis* |
| Gentianales | Gentianaceae | *Sabatia* | *Sabatia angularis* |

| | | | |
|---|---|---|---|
| Gentianales | Gentianaceae | *Sabatia* | *Sabatia bartramii* |
| Gentianales | Gentianaceae | *Sabatia* | *Sabatia brevifolia* |
| Gentianales | Gentianaceae | *Sabatia* | *Sabatia calycina* |
| Gentianales | Gentianaceae | *Sabatia* | *Sabatia campanulata* |
| Gentianales | Gentianaceae | *Sabatia* | *Sabatia difformis* |
| Gentianales | Gentianaceae | *Sabatia* | *Sabatia dodecandra* |
| Gentianales | Gentianaceae | *Sabatia* | *Sabatia grandiflora* |
| Gentianales | Gentianaceae | *Sabatia* | *Sabatia stellaris* |
| Liliales | Colchicaceae | *Uvularia* | *Uvularia grandiflora* |
| Liliales | Colchicaceae | *Uvularia* | *Uvularia perfoliata* |
| Liliales | Colchicaceae | *Uvularia* | *Uvularia puberula* |
| Liliales | Colchicaceae | *Uvularia* | *Uvularia sessilifolia* |
| Liliales | Liliaceae | *Clintonia* | *Clintonia borealis* |
| Liliales | Liliaceae | *Clintonia* | *Clintonia umbellulata* |
| Liliales | Liliaceae | *Erythronium* | *Erythronium americanum* |
| Liliales | Liliaceae | *Erythronium* | *Erythronium umbilicatum* |
| Liliales | Liliaceae | *Lilium* | *Lilium canadense* |
| Liliales | Liliaceae | *Lilium* | *Lilium catesbaei* |
| Liliales | Liliaceae | *Lilium* | *Lilium philadelphicum* |
| Liliales | Liliaceae | *Lilium* | *Lilium superbum* |
| Liliales | Melanthiaceae | *Trillium* | *Trillium catesbaei* |
| Liliales | Melanthiaceae | *Trillium* | *Trillium cernuum* |
| Liliales | Melanthiaceae | *Trillium* | *Trillium cuneatum* |
| Liliales | Melanthiaceae | *Trillium* | *Trillium erectum* |
| Liliales | Melanthiaceae | *Trillium* | *Trillium grandiflorum* |
| Liliales | Melanthiaceae | *Trillium* | *Trillium maculatum* |
| Liliales | Melanthiaceae | *Trillium* | *Trillium rugelii* |
| Liliales | Melanthiaceae | *Trillium* | *Trillium undulatum* |
| Liliales | Melanthiaceae | *Trillium* | *Trillium vaseyi* |
| Magnoliales | Annonaceae | *Asimina* | *Asimina angustifolia* |
| Magnoliales | Annonaceae | *Asimina* | *Asimina parviflora* |
| Magnoliales | Annonaceae | *Asimina* | *Asimina triloba* |
| Malpighiales | Passifloraceae | *Passiflora* | *Passiflora incarnata* |
| Malpighiales | Passifloraceae | *Passiflora* | *Passiflora lutea* |
| Malvales | Malvaceae | *Hibiscus* | *Hibiscus laevis* |
| Malvales | Malvaceae | *Hibiscus* | *Hibiscus moscheutos* |
| Malvales | Malvaceae | *Hibiscus* | *Hibiscus syriacus* |
| Malvales | Malvaceae | *Hibiscus* | *Hibiscus trionum* |
| Piperales | Aristolochiaceae | *Aristolochia* | *Aristolochia macrophylla* |
| Piperales | Aristolochiaceae | *Aristolochia* | *Aristolochia serpentaria* |
| Piperales | Aristolochiaceae | *Hexastylis* | *Hexastylis arifolia* |
| Piperales | Aristolochiaceae | *Hexastylis* | *Hexastylis virginica* |
| Rosales | Rosaceae | *Rosa* | *Rosa blanda* |
| Rosales | Rosaceae | *Rosa* | *Rosa carolina* |
| Rosales | Rosaceae | *Rosa* | *Rosa nitida* |
| Rosales | Rosaceae | *Rosa* | *Rosa palustris* |
| Rosales | Rosaceae | *Rosa* | *Rosa virginiana* |
| Solanales | Solanaceae | *Physalis* | *Physalis heterophylla* |
| Solanales | Solanaceae | *Physalis* | *Physalis longifolia* |
| Solanales | Solanaceae | *Physalis* | *Physalis pubescens* |
| Solanales | Solanaceae | *Physalis* | *Physalis walteri* |
